# Genetic diversity patterns of human ethnic groups as inferred from the 1000 genomes

**DOI:** 10.1101/2021.12.14.472684

**Authors:** Zhiyi Xia, Shi Huang

## Abstract

Human genetic diversity remains to be better understood. We here analyzed data from the 1000 Genomes Project and defined group specific fixed alleles (GSFAs) as those that are likely fixed in one ethnic group but non-fixed in at least one other group. The fraction of derived alleles in GSFAs indicates relative distance to apes because such alleles are absent in apes. Our results show that different groups differed in GSFA numbers consistent with known genetic diversity patterns, but also differed in the fraction of derived alleles in GSFAs throughout the entire genome, with East Asians having the largest fraction, followed by South Asians, Europeans, Native Americans, and Africans. Fast evolving sites such as intergenic regions were enriched with derived alleles and showed greater differences in GSFA numbers between East Asians and Africans. Furthermore, GSFAs in East Asians are mostly not fixed in other groups especially Africans, which was particularly more pronounced for fast evolving noncoding variants, while GSFAs in Africans are mostly also fixed in East Asians. Finally, variants that are likely non-neutral such as those leading to stop codon gain/loss and splice donor/acceptor gain/loss showed patterns similar to those of fast-evolving noncoding variants. These results can be accounted for by the maximum genetic diversity theory but not by the neutral theory or its inference that Eurasians suffered bottlenecks, and have implications for better management of group specific genetic diseases.

## Introduction

Humans and great apes are believed to share a common ancestor. Nucleotide difference between the human and chimpanzee genomes is remarkably small, representing only about 2% of the genome, but presumably underlies the traits that make human unique or distinct from apes (Prado-Martinez et al., 2013). Modern humans are more differentiated from the ancestral state than the archaic humans in terms of both genotypes and phenotypes, even though they had coexisted and interbred with each other at ~50000 years ago as indicated by the fossil record and ancient DNAs (Green et al., 2010; Meyer et al., 2012). Relative to modern humans, Neanderthals have more ancestral alleles or less derived alleles and more ancestral traits such as large brow ridges (Coon, 1962; Green et al., 2010; Wolpoff and Caspari, 2007).

Genetic mutations introduce into human populations single nucleotide polymorphisms (SNPs) that make two different human individuals differ in ~0.1% of their genome sequence. If the first modern human individual had 2% non-identity in DNA from chimpanzee, his/her descendants may incur new mutations in the 2% non-identical region that would represent back mutations to the ancestral state. The descendants may also sustain new mutations in the 98% identical region. These polymorphisms as introduced by mutations, together with variants introduced by admixture with less modern humans, would make human individuals non-identical in their sequence similarities with apes.

While different human ethnic groups are thought to be highly alike, they do show measurable genetic differences measured as averages, which may underlie certain trait differences such as susceptibility to certain diseases (Carrot-Zhang et al., 2021; Oni-Orisan et al., 2021). They are thought to show different degrees of admixture with archaic humans (Green et al., 2010; Meyer et al., 2012). They also show different genetic diversity levels, with Africans the highest and East Asians the lowest (Auton et al., 2015). Africans are further known to carry more ancestral alleles in both uniparental DNAs and autosomes as indicated by the rooting in Africa in phylogenetic trees rooted by the outgroup rooting method, which places the root in the human group that shares the most sequence identity with the outgroup chimpanzee (Ingman et al., 2000; Li et al., 2008; Underhill et al., 2000).

Genetic diversity patterns among living species are highly complex, which has long remained a challenge to molecular evolutionary theories (Huang, 2016; Kern and Hahn, 2018; Lewontin, 1974). The popular theory used for explaining genetic diversity patterns is the neutral theory, which postulates that genetic diversity level is positively related to the effective population size Ne (Kimura, 1983). However, Ne cannot be directly measured and is often tautologically derived by using the very genetic variables that it is meant to explain or predict. Ne is presumably proportional to census population size. However, the census population size of great apes is many orders of magnitude smaller than that of humans but the within species genetic diversity levels of great apes are much higher than that of humans (Prado-Martinez et al., 2013). This is explained away by the neutral theory by postulating an ad hoc solution, i.e., bottlenecks in humans, even though humans show no evidence of more inbreeding or more runs of homozygosity than apes (Prado-Martinez et al., 2013). Similarly, the low genetic diversity of Eurasians relative to Africans is also explained by ad hoc bottlenecks in Eurasians as in the Out of Africa model of modern human origins (Scerri et al., 2018). The assumption of neutrality for most standing variants by the neutral theory also remains uncertain or unproven. Rapid progress in functional genomics in the past two decades are steadily increasing the proportion of functional regions in the genome, especially in regions commonly thought as neutral (Dunham et al., 2012; Pouyet et al., 2018; Press et al., 2018; Quinodoz et al., 2021; Tsuzuki et al., 2020). Thus, models derived from the controversial neutral theory such as the Out of Africa model or the bottlenecks in Eurasians remain at best uncertain.

Many in the past have tried to use natural selection rather than neutrality to explain genetic diversity patterns (Hahn, 2008; Lewontin, 1974). The more recent theoretical development is the maximum genetic diversity (MGD) theory (Hu et al., 2013; Huang, 2008, 2009, 2016). By introducing the novel concept of physiological selection and maximum genetic distance or diversity as restricted by physiological complexity, MGD theory has coherently explained major genetic diversity patterns. Thus, that humans have lower genetic diversity than apes is because humans, being more complex, require building parts of higher precision, and hence lower level of random errors in its genome. Mutations that cause diseases or cannot be tolerated as polymorphisms in humans are often normal variations in animals (Kondrashov et al., 2002). The MGD of humans must be smaller than that of apes. Genetic diversities in humans in most genomic regions are already today at maximum or optimum levels as shown by higher genetic diversity in patient groups relative to controls (Chen et al., 2020; Gui et al., 2017; He et al., 2017; Lei et al., 2018; Zhu et al., 2015), and by the more sharing of fast evolving variants among ethnic groups due to convergent mutations (Yuan et al., 2019). MGD theory considers most variants to be functional or under selection, particularly under physiological selection. Fast evolving DNAs are more adaptive than slowly evolving DNAs and so under more selection (Kasinathan et al., 2020; Wang et al., 2020). A mutation deleterious to the physiology of the organism would not be able to make a normal healthy individual.

We here further investigated the pattern of human genetic diversity throughout the whole genome by using the previously published data of the 1000 genomes (1kG) project (phase 3) that include five ethnic groups. We uncovered remarkable groups specific features and differences among ethnic groups with regard to differentiation from the chimpanzee genome. The results cannot be explained by bottlenecks or the neutral theory but are more consistent with the MGD theory, and may help better understanding the implications of group specific health issues.

## Results

If bottlenecks were the reason for the low genetic diversity in East Asians, regions of the genome thought to have evolved at different rates should show similar degree of reduction in genetic diversity. A population with low genetic diversity should show higher numbers of fixed alleles. We downloaded and studied the Variant Call Format (VCF) file of the 1kG project (phase 3) that has annotated variants in terms of allele frequency in different human groups and of location in different types of DNAs (Auton et al., 2015). The 1kG project contains five ethnic groups, AFR (Africans), AMR (native Americans), EUR (Europeans), SAS (South Asians), and EAS (East Asians). We defined group specific fixed alleles (GSFAs) as those alleles that are different from the reference alleles and are fixed in one ethnic group but are non-fixed in at least one other group. In other words, GSFAs are alternative (Alt) alleles and have alt allele frequency 1 (AF=1) in one group but have AF<1 in at least one other group. We also defined non-group specific fixed alleles (NGSFAs) as those alleles that are the same as the reference alleles and are fixed in one ethnic group but are non-fixed in at least one other group. Specifically, NGSFAs are reference alleles and have alt allele frequency 0 (AF=0) in one group but have AF>0 in at least one other group. NGSFAs are present in at least two different set of individuals, one represented by the individuals serving as the reference genome and the other represented by one of the 5 groups in 1kG. Therefore, NGSFAs are more commonly shared among ethnic groups whereas GSFAs are more group specific.

Groups with low genetic diversity should have more GSFAs and NGSFAs relative to groups with higher genetic diversity. If the low genetic diversity of EAS was due to bottlenecks as theorized by the neutral theory, one would expect similar degree of increase in GSFA/NGSFA numbers across all types of variants irrespective of their evolutionary rates. We examined the following types of DNA sequences that are known to differ in evolutionary rates, with intergenic variants having the highest evolutionary rates, followed by intron variants, synonymous variants, missense variants, and slow missense variants. The slow missense variants here are a subset of missense variants found in the slowest evolving set of proteins that has been defined previously and have been shown to be the most neutral (Wang et al., 2020; Yuan et al., 2019). In addition, we also examined stop codon gain or loss variants and splice donor or acceptor gain or loss variants, which affect protein structure and function more severely and are hence likely to be non-neutral. We enumerated GSFA and NGSFA numbers for each of the five groups in 1kG for each type of variants and calculated the ratio of AFR versus EAS in GSFA or NGSFA numbers. As shown in Figure 1, the AFR/EAS ratio was <1 for all types of variants examined, consistent with the known lower genetic diversity in EAS relative to AFR. Remarkably, the ratio differed based on the evolutionary rates of variants, with the fastest evolving variants showing the lowest ratio and the slow set of missense variants the highest. Furthermore, the ratio was lower in GSFA relative to NGSFA, indicating dramatic difference between GSFA and NGSFA. These observations are unexpected from the bottleneck hypothesis. The ratio for stop/splice variants was between that for the missense variants and the slow set of missense variants. Since the results here showed lower ratios in missense variants compared to the slowest evolving missense variants, one would expect other non-neutral variants to show the same. Indeed, stop/splice variants also showed lower ratio compared to the slow set and are hence less neutral than the slowly evolving missense variants. Also consistently, EAS showed lower number of common SNPs (MAF>=0.05) than AFR, less so in the slow set of missense variants, and the stop/splice variants again were more like the fast evolving noncoding variants and the missense variants (Figure 2). Given that the noncoding variants also showed lower ratios than the neutral slow set of missense variants, one can infer that the noncoding variants are non-neutral. The faster evolutionary rates of non-coding variants may explain why they appear to be the least neutral among the variants examined here (only fast mutation can meet fast adaptive needs).

**Figure 1.**
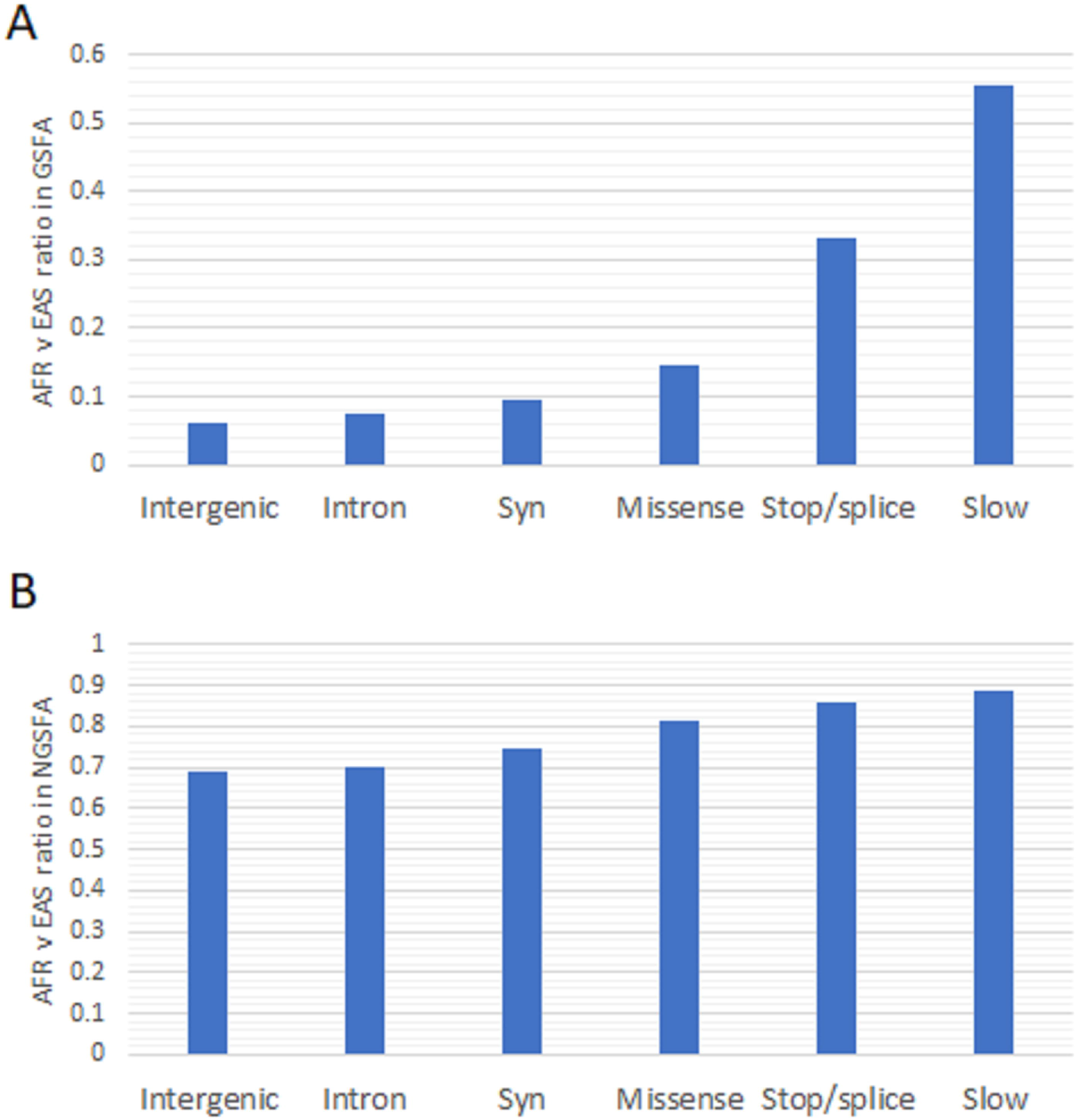
Ratio of AFR vs EAS in GSFA and NGSFA numbers. GSFA and NGSFA numbers were counted for each type of DNA as shown for AFR and EAS and the ratio of AFR vs EAS in GSFA (A) and NGSFA numbers (B) are shown.

**Figure 2.**
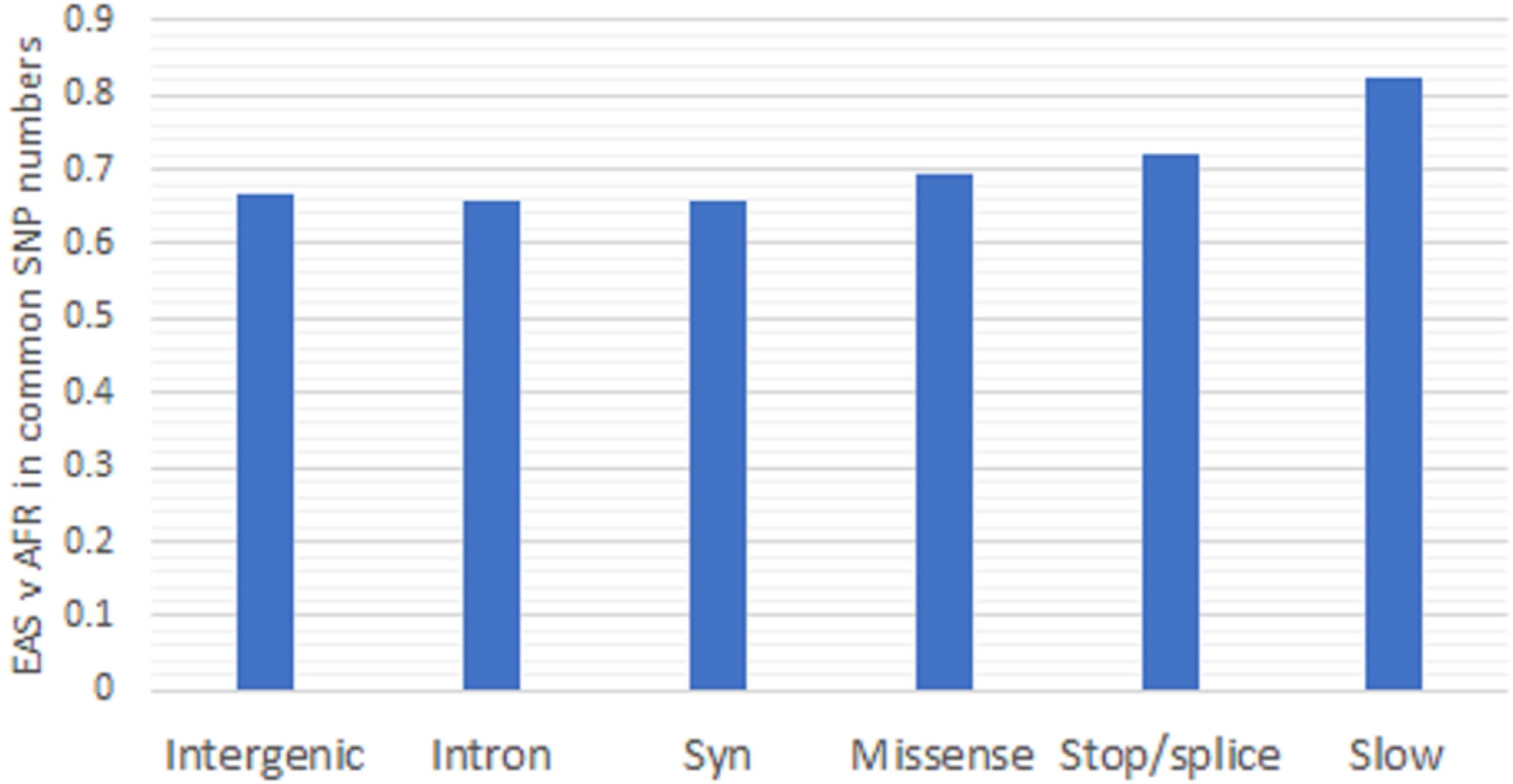
Ratio of EAS vs AFR in common SNP numbers. Common SNPs are defined as having MAF>=0.05.

An ethnic group with low levels of genetic diversity, and hence high numbers of GSFA or NGSFA, may not differ in relative distance to apes from those with higher levels as the fraction of derived alleles (different from chimpanzee allele) in GSFA or NGSFA may not differ, if low genetic diversity was due to bottlenecks which should affect both derived and ancestral alleles similarly. We next assigned derived allele status to the alt allele by comparing to the chimpanzee reference genome and calculated the fraction of derived alleles in GSFA or NGSFA. As shown in Figure 3, higher fraction of derived alleles was found in GSFA than in NGSFA, indicating enrichment of derived alleles in group specific variants. Fast evolving variants showed higher fraction of derived alleles relative to slowly evolving variants. GSFAs in the slow set of missense variants and in stop/splice variants showed no derived alleles for this analysis to be informative in GSFAs. Different ethnic groups showed different fraction of derived alleles with EAS the highest, followed by SAS, EUR, AMR and AFR. The data suggests that the EAS genomes may be the most differentiated from the chimpanzee genome.

**Figure 3.**
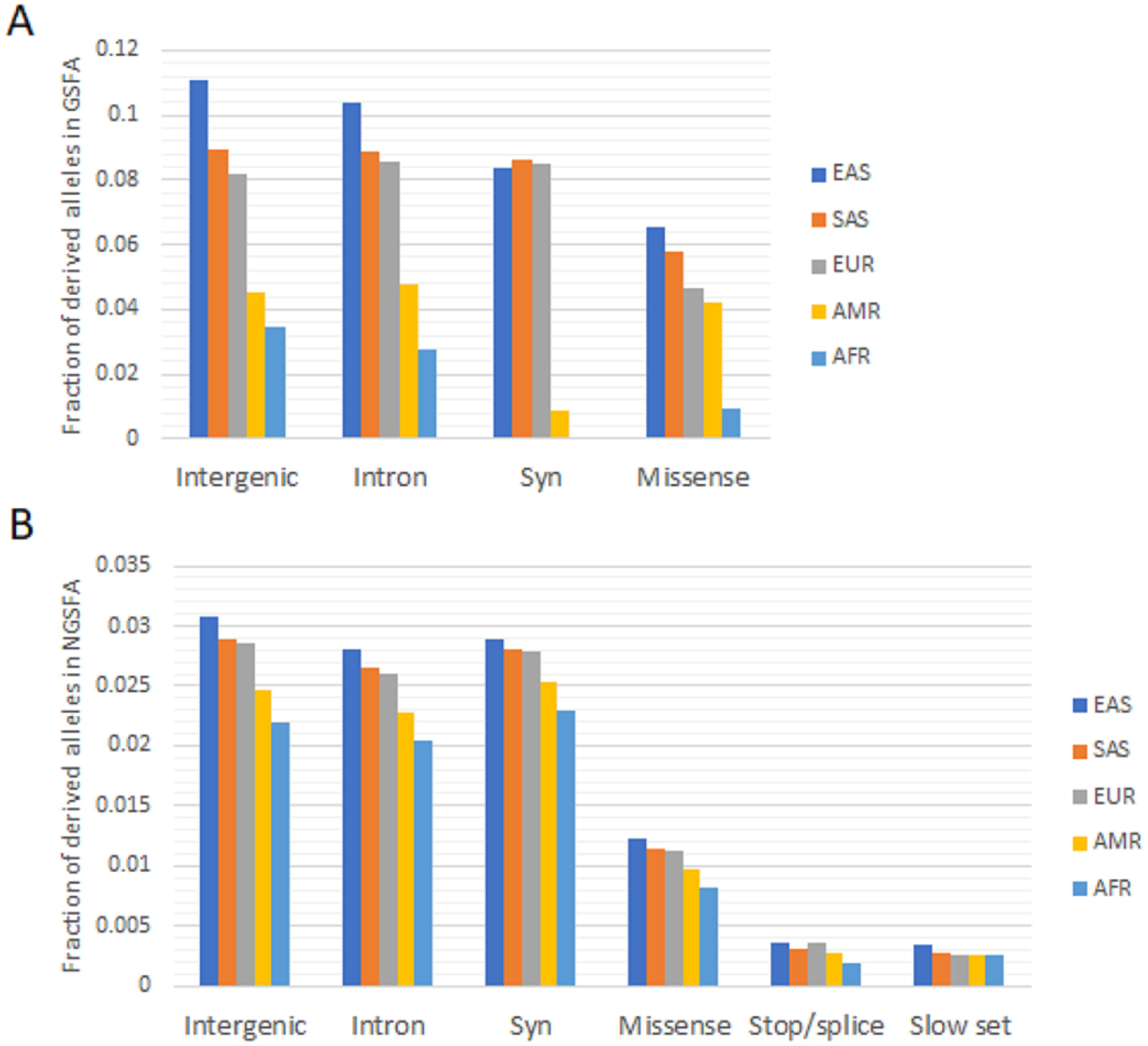
Fraction of derived alleles in fixed alleles. Fraction of derived alleles in GSFAs (A) and NGSFAs (B) are shown for each ethnic group.

We next studied among GSFAs or NGSFAs in EAS how many are non-fixed in AFR, and vice versa. The results showed that most GSFAs in EAS were not fixed or were polymorphic in AFR (MAF>0), particularly so for fast evolving variants (Figure 4A). In contrast, most GSFA in AFR were also fixed in EAS, irrespective of the evolutionary rates of the variants. Results on NGSFAs were similar but less dramatic. These observations further indicate remarkable differences in GSFAs among different ethnic groups.

**Figure 4.**
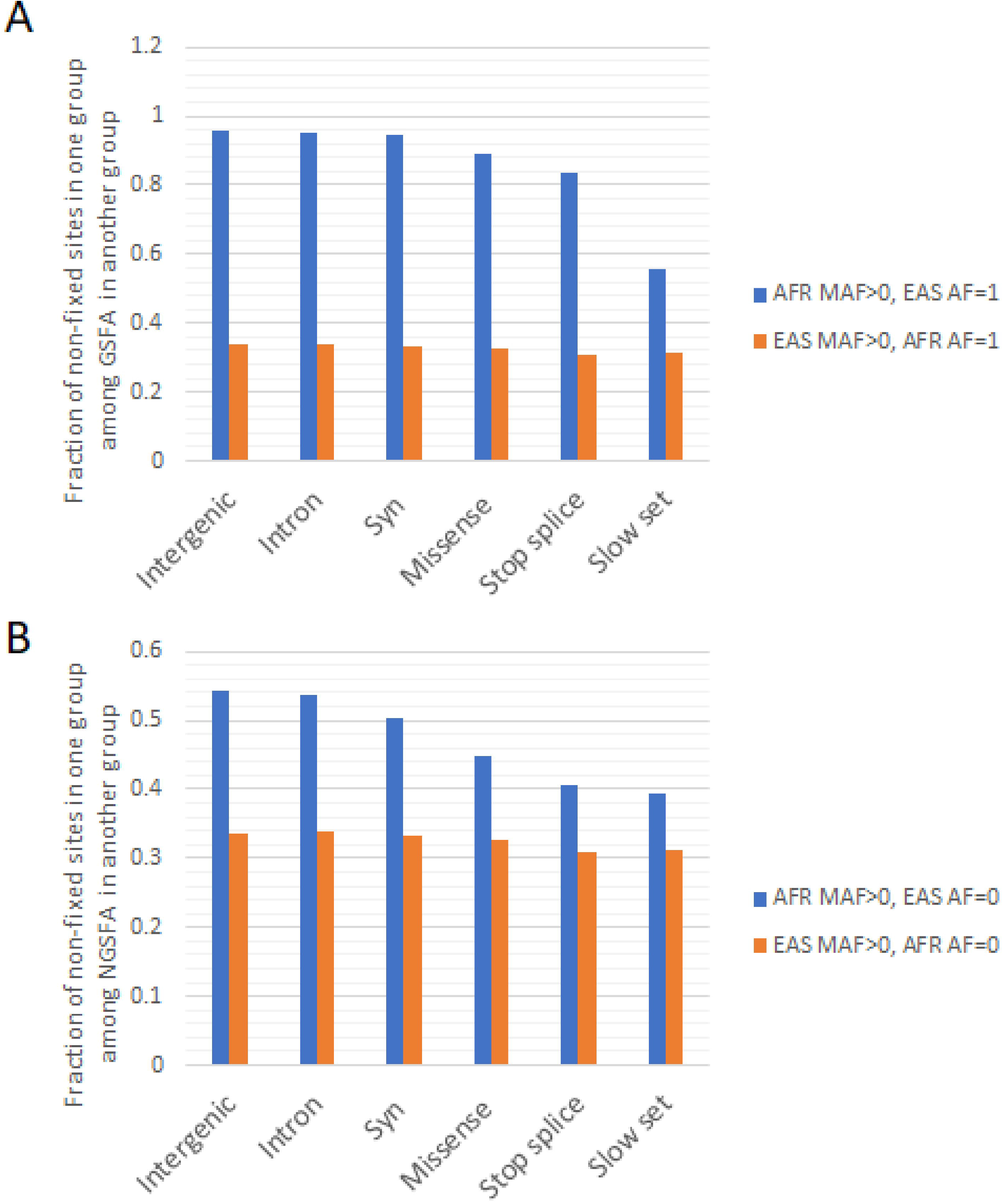
Fraction of non-fixed sites. For each type of DNA, shown are fractions of non-fixed sites (MAF>0) in one group among fixed alleles (AF=1 or 0) in another group. **A.** Fraction of non-fixed sites in GSFAs. **B.** Fraction of non-fixed sites in NGSFAs.

## Discussions

Our study here uncovered novel genetic diversity patterns of humans. Variants in fast evolving intergenic DNAs showed distinct patterns from that of the slowly evolving set of missense variants. Which type of variants are neutral? The observations here suggest that the slowly evolving set of missense variants are neutral whereas the fast evolving intergenic DNAs are functional, which is consistent with previous studies (Wang et al., 2020; Yuan et al., 2019). Slowly evolving missense variants are more neutral than fast evolving missense variants or have more conservative amino acid changes (Wang et al., 2020). The results here showed that fast evolving missense variants showed lower ratio of AFR vs EAS in GSFAs compared to the slowly evolving missense variants. Thus, functional variants should show lower ratio of AFR vs EAS in GSFAs compared to the neutral variants. Since variants in intergenic DNAs showed the lowest ratio of AFR vs EAS in GSFAs, they are likely to be functional. Relative to the slowly evolving set of missense variants, missense variants as a whole (including the slow set) also showed reduced ratio of EAS vs AFR in common SNP numbers, and higher fraction of derived alleles in NGSFAs. So also did the intergenic variants, again indicating that they are likely to be functional. Furthermore, variants that change stop codons or splice site donors and acceptors are presumably functional and indeed showed patterns more like the missense variants as a whole. Overall, the faster evolving the variants, the less neutral, which is consistent with the intuitive expectation that only fast evolving mutations could meet the adaptive needs of fast changing environments. Indeed, the dogma that less conserved DNAs are less functional has been overturned (Kasinathan et al., 2020; Wang et al., 2020).

Several observations here were inconsistent with the neutral theory’s bottleneck hypothesis to explain the low genetic diversity of East Asians. First, we found that the relatively low genetic diversity as measured here by GSFAs and NGSFAs in East Asians varied depending on the evolutionary rates of the types of DNAs concerned. Fast evolving DNAs such as intergenic DNAs showed more pronounced differences in genetic diversity between ethnic groups. However, the bottleneck hypothesis expects reduced genetic diversity levels irrespective of evolutionary rates. Second, GSFAs showed more dramatic reduction of genetic diversity in EAS relative to AFR than NGSFA did. Third, the fraction of derived alleles in GSFAs also varied depending on the evolutionary rates of the nucleotides concerned. Fourth, we found higher fraction of derived alleles in GSFAs in East Asians relative to that in Africans. However, the bottleneck hypothesis expects no changes in the fraction of derived alleles in the group suffering the bottleneck relative to the group without the bottleneck. Finally, we found that alleles fixed in East Asians are mostly not fixed in Africans, especially for fast evolving DNAs such as intergenic DNAs, but alleles fixed in Africans are mostly also fixed in East Asians. While the bottleneck hypothesis does expect the group suffering a bottleneck to have fixed alleles that would not be fixed in the ancestral group without the bottleneck, it however cannot explain the association of this fixation with the evolutionary rates of the variants. There are also other studies not favoring the bottleneck hypothesis, where the length of long runs of homozygosity (ROH) relative to the length of short ROH was found to be similar for Africans and Eurasians and much lower than that in people from Middle East who are known to practice consanguineous marriages (Scott et al., 2016).

The nearly neutral theory of Ohta suggests that slightly deleterious variants should behave like neutral ones in small populations as a result of bottlenecks (Ohta, 1973). Thus, variants affecting stop/splice sites are expected to accumulate in EAS, leading to lower number of fixed alleles in EAS. This may explain why stop/splice variants are more like the neutral variants as represented by the slowly evolving set of missense variants. However, this nearly neutral perspective cannot explain the difference among supposedly neutral variants, such as the difference in the numbers of GSFAs and NGSFAs among intergenic, intron, and syn variants. It also cannot explain the different fractions of derived alleles among ethnic groups.

Our results here appear to be well expected by the MGD theory. Most DNAs are functional rather than neutral. Fast evolving DNAs such as noncoding intergenic DNAs play more adaptive roles in response to fast changing environments and are hence under more natural selection. During human evolution, the human physiology becomes more complex especially for the nervous system. This must require an increase in the precision of the genome and so a reduction in MGD or in the maximum amounts of random errors allowed. This reduction would most affect the fast evolving functional DNAs and least affect the slowly evolving neutral DNAs. Variants not allowed in groups with lower MGD could be allowed in groups with higher MGD, especially for fast evolving functional variants that are under more selection. Variants not allowed (selected against) in groups with higher MGD would also be not allowed in groups with lower MGD, irrespective of evolutionary rates of the variants. This has implications for the identification of disease risk mutations among ethnic groups (Carrot-Zhang et al., 2021; Oni-Orisan et al., 2021). Future studies may uncover those types of group specific disease risk mutations.

## Methods

We downloaded the VCF file of the 1kG project (phase 3) from the 1kG project website (Auton et al., 2015). The file annotated each SNP according to DNA types, such as intergenic, intron, synonymous, missense, stop codon gain/loss, and splice donor/acceptor gain/loss. It also has alt allele frequency data. From the missense SNPs, we picked out the slowly evolving set of SNPs that are missense variants found in the slowest evolving set of proteins as defined previously (Wang et al., 2020; Yuan et al., 2019). We removed noninformative SNPs that have AF=1 or AF=0 across all five ethnic groups. The allele frequency data of each SNP was taken directly from the VCF file and analyzed for enumerating GSFAs and NGSFAs. GSFAs are group specific fixed variants that show AF=1 in one group and AF<1 in at least one other group. NGSFA are non GSFAS that show AF=0 in one group and AF>0 in at least one other group.

### Assign derived allele status

Ancestral status means identical to the chimpanzee sequence, and derived status means different. Chimp (Clint_PTRv2/panTro6) reference genome sequence was downloaded from UCSC genome browser (http://hgdownload.soe.ucsc.edu/goldenPath/panTro6/bigZips/). To match human genome reference (hg19) coordinates with chimpanzee reference genome coordinates (panTro6), we downloaded the chain file hg19ToPanTro6.over.chain from UCSC genome browser (http://hgdownload.cse.ucsc.edu/gbdb/hg19/liftOver/). We then used GATK-4.1.8.1(https://github.com/broadinstitute/gatk/releases) LiftOverIntervalList to convert hg19 coordinates into chimp(panTro6) coordinates (Kuhn et al., 2012). We next used GATK-4.1.8.1 ExtractSequences to obtain the genotype of panTro6 genome based on the chimp coordinates (Van der Auwera et al., 2013). If neither allele of a variant matched the chimpanzee allele, the variant was noninformative and excluded from analysis.

## Acknowledgments

Supported by the National Natural Science Foundation of China grant 81171880 (S.H.).

## Additional Information

### Author contributions

Z.X. performed data analysis. S.H. devised the project, analyzed the data, and wrote the manuscript. All authors edited the manuscript.

